# Hyperinsulinemia promotes HMGB1 release leading to inflammation induced systemic insulin resistance: An interplay between pancreatic beta-cell and peripheral organs

**DOI:** 10.1101/705103

**Authors:** Abhinav Choubey, Aditya K Kar, Khyati Girdhar, Tandrika Chattopadhyay, Surbhi Dogra, Shaivya Kushwaha, Bikash Medhi, Anil Bhansali, Chinmay Kumar Mantri, Ullas Kolthur-Seetharam, Debabrata Ghosh, Prosenjit Mondal

## Abstract

Insulin resistance results from several pathophysiologic mechanisms, including chronic tissue inflammation and defective insulin signaling. Pancreatic β-cells hypersecretion (hyperinsulinemia), is a central hallmark of peripheral insulin resistance. However, the underlying mechanism by which hyperinsulinemia perpetuates towards the development of insulin resistance remains unclear and is still a bigger therapeutic challenge. Here, we found hyperinsulinemia triggers inflammation and insulin resistance by stimulating TLR4-driven inflammatory cascades. We show that hyperinsulinemia activates the TLR4 signaling through HMGB1, an endogenous TLR4 ligand emanating from hyperinsulinemia exposed immune cells and peripheral organs like adipose tissue and liver. Further, our observation suggests hyperinsulinemia ensuring hyperacetylation, nuclear-to-cytoplasmic shuttling and release of HMGB1 into the extracellular space. HMGB1 was also found to be elevated in serum of T2DM patients. We found that extracellular HMGB1 plays a crucial role to promote proinflammatory responses and provokes systemic insulin resistance. Importantly, *in-vitro* and *in-vivo* treatment with naltrexone, a TLR4 antagonist led to an anti-inflammatory phenotype with protection from hyperinsulinemia mediated insulin resistance. *In-vitro* treatment with naltrexone directly enhanced SIRT1 activity, blocked the release of HMGB1 into extracellular milieu, suppressed release of proinflammatory cytokines and ultimately led to insulin-sensitizing effects. These observations elucidate a regulatory network between pancreatic β-cells, macrophage and hepatocytes and assign an unexpected role of TLR4 - HMGB1 signaling axis in hyperinsulinemia mediated systemic insulin resistance.

**Graphical Abstract:** 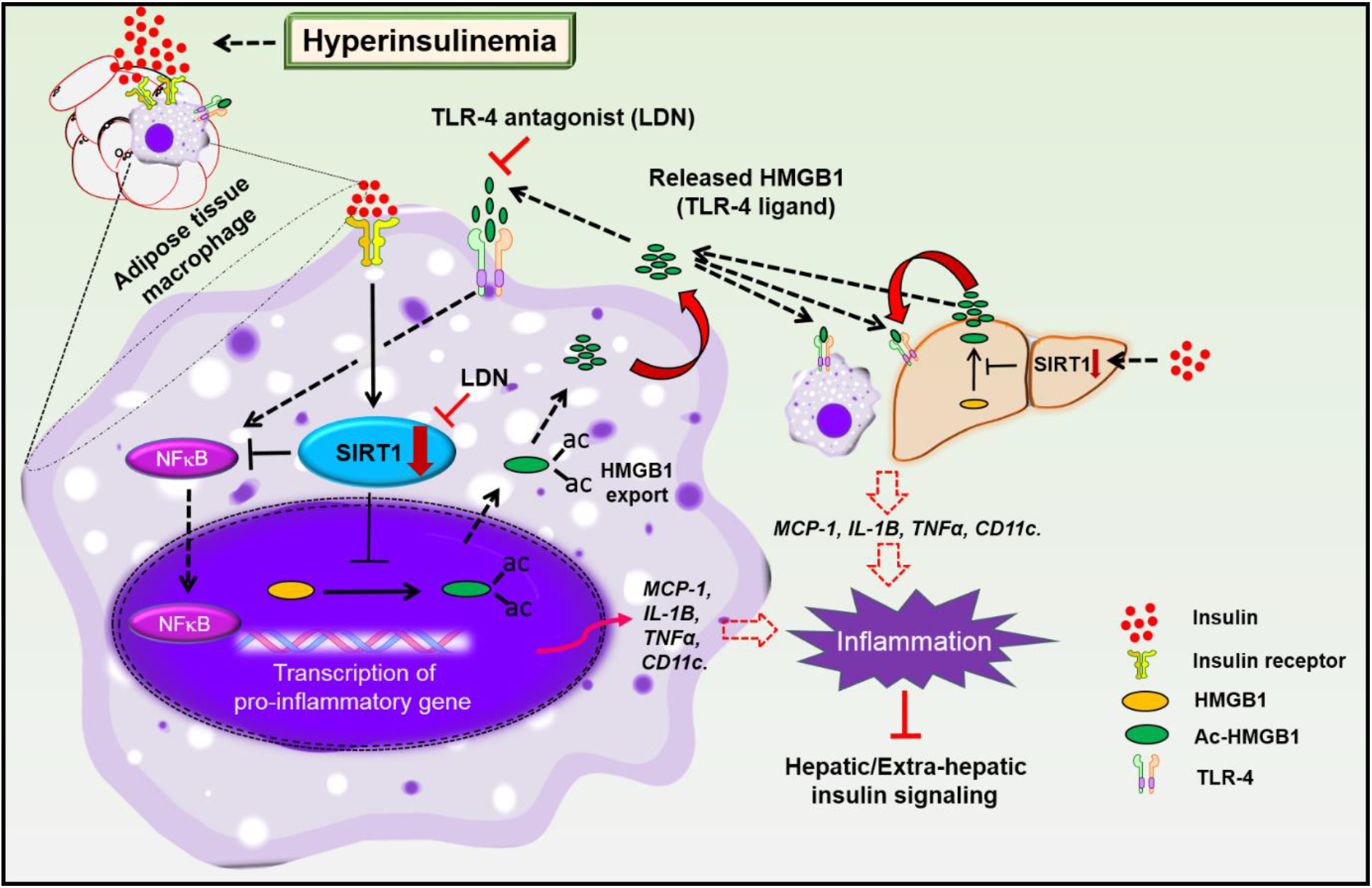

## Introduction

The incidence of diabetes, obesity and Non-Alcoholic Fatty Liver Diseases (NAFLD) has reached epidemic status in the World. Insulin resistance is a common link in the development of these conditions (Garvey et al., 1988; Olefsky et al., 1988; Reaven, 2005). Insulin resistance, a complex pathophysiologic event characterized by defective insulin signaling in metabolically active tissues. Free fatty acids and soluble inflammatory mediators released from immune cells and peripheral organs (adipose tissue, skeletal muscle, Liver etc) are contributed in regulating insulin action (Weisberg et al., 2003; Xu et al., 2003). Adipocytes release adipokines (adiponectin, leptin etc) and proinflammatory cytokines, including tumor necrosis factor-a (TNFa) interleukin-6 (IL-6), IL8, etc are recognized to contribute insulin resistance (Blüher, 2012)(Blüher and Mantzoros, 2015). SIRT1, a mammalian ortholog of Sir2 (silent information regulator 2), and type III NAD^+^-dependent deacetylase, is reported to repress inflammatory responses. Downregulation of SIRT1 in adipose tissue induced proinflammatory phenotype via NF-kB driven inflammation (Yoshizaki et al., 2009). Whereas, overexpression of SIRT1 prevents high-fat diet-induced adipose tissue inflammation (Pfluger et al., 2008). Down regulation of SIRT1 level also ensures hyperacetylation and release of HMGB1; one of the key proteins regulating inflammatory response (Rickenbacher et al., 2014). HMGB1 is a DNA binding protein and involves in transcriptional regulation and nucleosome stabilization. In addition to its nuclear function, HMGB1 can have crucial extracellular role as an inflammatory cytokine (Klune et al., 2008; Thomas, 2001).

Liver also participates in causing defective insulin signaling by releasing proteins into the circulation; these liver derived proteins are termed as hepatokines (Stefan and Häring, 2013). The hepatokine fetuin-A was identified as an endogenous ligand for TLR4 through which saturated fatty acids induce proinflammatory cascade and regulating insulin action (Pal et al., 2012). Other than fetuin-A, hepatokines Fibroblast growth factor 21 (FGF21) and Selenoprotein P are major drivers of insulin resistance (Badman et al., 2007; Misu et al., 2010).

Like adipose tissue and liver, pancreatic β-cells can also contribute to insulin resistance. Hypersecretion of β-cell is considered to be a compensatory response to insulin resistance. However, a growing number of evidences have suggested, hyperinsulinemia occurs prior to development of hepatic and central insulin resistance and may actually drive and/or perpetuate towards insulin resistance. However, the mechanism by which elevated hyperinsulinemia derails insulin signaling remains unclear.

Pathologically high insulin level has emerged as an early cause of insulin resistance and other metabolic diseases like NAFLD, obesity and T2DM (Chen et al., 2018; Rhee et al., 2011; Steneberg et al., 2015; Sung et al., 2011). Moreover, increase in plasma insulin levels in rodents have been shown to induce insulin resistance resembling clinical state (Marín-Juez et al., 2014; Pereira et al., 2014; Shoelson et al., 2006; Wang et al., 2016). Increased circulating levels of insulin have been shown to induce NAFLD, causing insulin resistance mainly by stimulating hepatic expression of the FA transporter Cd36 (Badman et al., 2007). Hyperinsulinemia was also seen to induce M1 phenotype in organ-resident macrophages, which are known to induce inflammatory signaling pathway (Su et al., 2016). However, understanding the role of hyperinsulinemia in the etiology of insulin resistance remains enigmatic; hence, identification of pathways that influence the pathogenesis of insulin resistance has the potential to lead new therapies for preventing or delaying onset of disease.

Using numerous complementary *in-vitro* and *in-vivo* experimental approaches we have demonstrated here how hyperinsulinemia increases circulatory levels of HMGB1; an endogenous TLR4 ligand and its possible links to the development of systemic insulin resistance. Here, we investigated the role of known TLR4 antagonist; naltrexone (Weerts et al., 2008), in the pathophysiology of hyperinsulinemia mediated insulin resistance. Our data suggests, treatment with the low dose naltrexone (LDN), suppresses hyperinsulinemia mediated HMGB1’s cytosolic and NF-kBp65 nuclear translocation mainly through SIRT1. Importantly, humans with T2DM also exhibit increased serum HMGB1 levels. Thus, providing a proof of concept that hyperinsulinemia induced systemic insulin resistance arises secondary to the inflammatory response caused by secretion of HMGB1 from immune cells and peripheral organs.

## Results

### Hyperinsulinemia induced proinflammatory cytokine expression requires TLR4-dependent inflammatory events

Macrophages, specifically proinflammatory M1 cells, have been associated with insulin resistance, obesity and diabetes (Weisberg et al., 2003). However, how insulin signaling in macrophages leads to systemic insulin resistance is poorly addressed. Towards this, we challenged macrophage cells with high dose of insulin (100nM) for 24 hrs in presence and absence of TLR4 antagonist (LDN). Insulin treatment induced expression of *IL*-1β, *MCP*-1, and *CD11c* which are associated with M1 macrophage phenotype and to our surprise this induction in M1 cytokine expression was attenuated by LDN (Fig-1A). No significant change in M2 cytokines was observed in either group. To further verify the role of TLR4 in modulating hyperinsulinemia mediated inflammation, macrophage cells were transfected with TLR4 specific siRNA and then treated with high insulin. As shown in (Fig-S1), high insulin treatment was unable to induce gene expression of proinflammatory markers in TLR4-deficient macrophage cells.

**Figure 1:**
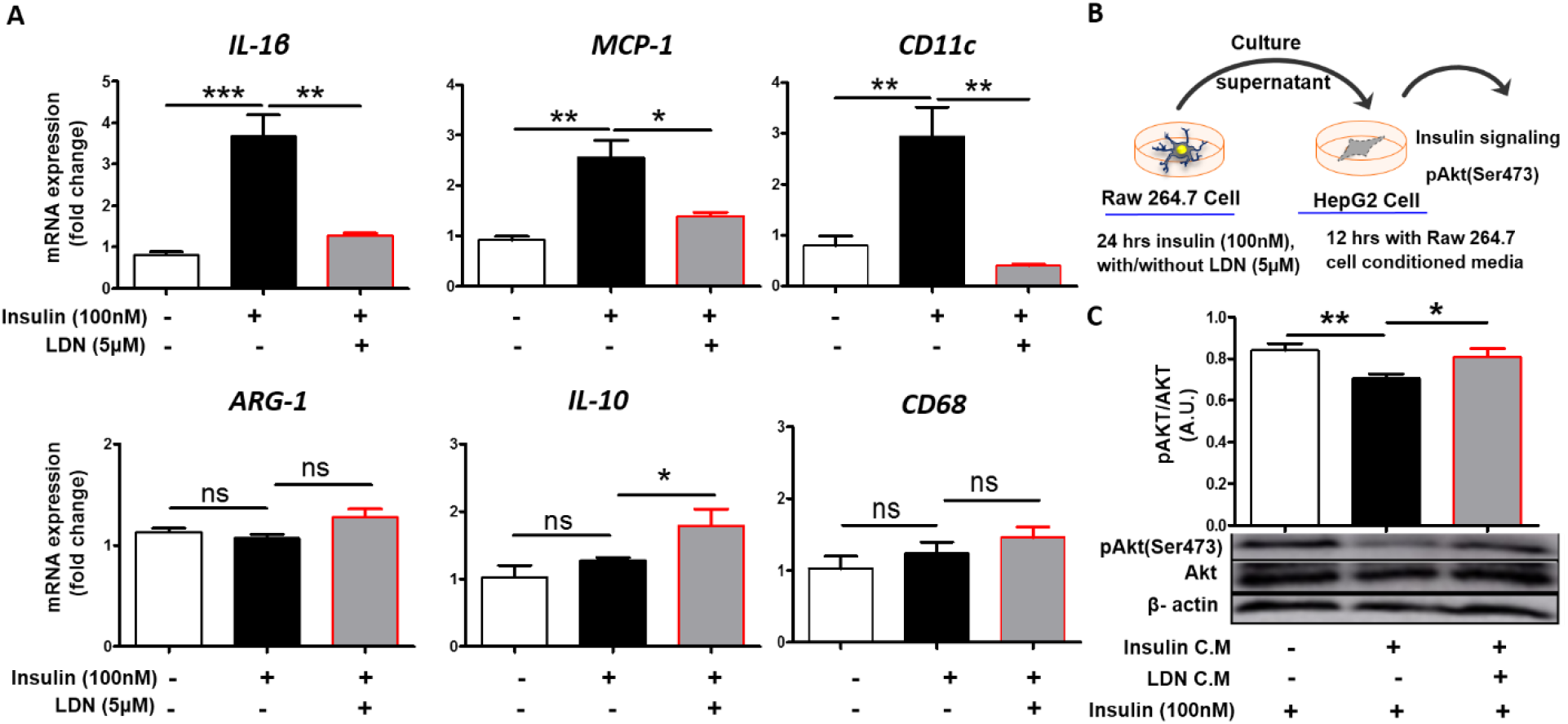
Hyperinsulinemia induces proinflammatory cytokine expression from macrophage cells and modulates the crosstalk between macrophage and liver: (**A**) qRT-PCR of indicated genes (*IL-1β, MCP-1, CD11c, ARG-1, IL-10, CD68*) in murine macrophage (Raw264.7). (**B**) Schematic depicting collection of conditioned media from macrophage cells and treatment to hepatic cells (HEPG2). Briefly, conditioned media from macrophage cells treated with 100nM insulin in presence or absence of LDN was collected and challenged (in half dilution with DMEM serum starved media) to HepG2 cells for 12 hours. (**C**) IB in HepG2 Cell lysates using anti pAkt (ser473), Akt and β-actin antibodies after indicated conditioned media treatment, Values are expressed as mean ± SEM (n=3) (mean ± SEM ***p<0.001, **p<0.01 *p<0.05.)

To check if the factors released by M1 macrophage can induce hepatic insulin resistance, we treated HepG2 cells with conditioned media from macrophage cells (Fig-1B), and interestingly found that conditioned media from hyperinsulinemic group blunted the phosphorylation of Akt (Ser473), measured here as a surrogate for insulin resistance, compared to conditioned media from untreated group. This increase in insulin resistance was abrogated in presence of LDN, indicating TLR4 is associated with hyperinsulinemia mediated insulin resistance (Fig-1C). To delineate, whether cytokines released by macrophage or any other soluble factor is responsible for inhibition of phosphorylation, we treated HepG2 cells with heat-inactivated hyperinsulinemia challenged macrophage conditioned media and found that heat inactivation of conditioned media did not blunt Akt (Ser473) phosphorylation in HepG2 cells (Fig-S2). Together these finding suggests that certain factor secreted by insulin-challenged macrophage cell confer hepatic insulin resistance in our *in-vitro* model which is inhibited by TLR4 antagonist.

### TLR4 antagonist prevents hyperinsulinemia induced ATM stimulated cytokines expression and insulin resistance in mice

We developed a short-term high fat diet (HFD) fed hyperinsulinemic mice model (Fig-2A), to provide a deeper understanding of endogenous mechanisms that regulate insulin action. As shown in (Fig-2B), fasting serum insulin level was higher in short term HFD-Saline fed mice than in NCD (normal chow diet) fed mice. Interestingly, fasting insulin levels were decreased significantly after LDN treatment in HFD-LDN group. Dynamic glucose tolerance tests (ipITT and ipGTT) were carried out to find out peripheral response to insulin and glucose as well as to explore *in-vivo* efficacy of LDN on glucose homeostasis and whole-body insulin sensitivity (Dogra et al., 2019). HFD fed mice displayed impairment in insulin sensitivity (measured by higher area under curve) as compared to NCD groups and the insulin sensitivity returned to the normal level when HFD fed mice were treated with LDN (Fig-2C). A similar trend was observed for ipGTT, wherein HFD fed mice exhibited an increased glucose excursion AUC as compared to mouse on NCD which was significantly reduced in HFD-LDN mice (Fig-2D). Together, these findings indicated the potential role of LDN in improving glucose tolerance and insulin sensitivity in diet induced hyperinsulinemic mice. Adipose tissue macrophage (ATM) is closely linked to this inflammatory condition which leads to insulin resistance (Blüher, 2012). Increased accumulation of proinflammatory or classically activated (M1) macrophage in ATM is positively correlated with, insulin resistance (Shin et al., 2017).

**Figure 2:**
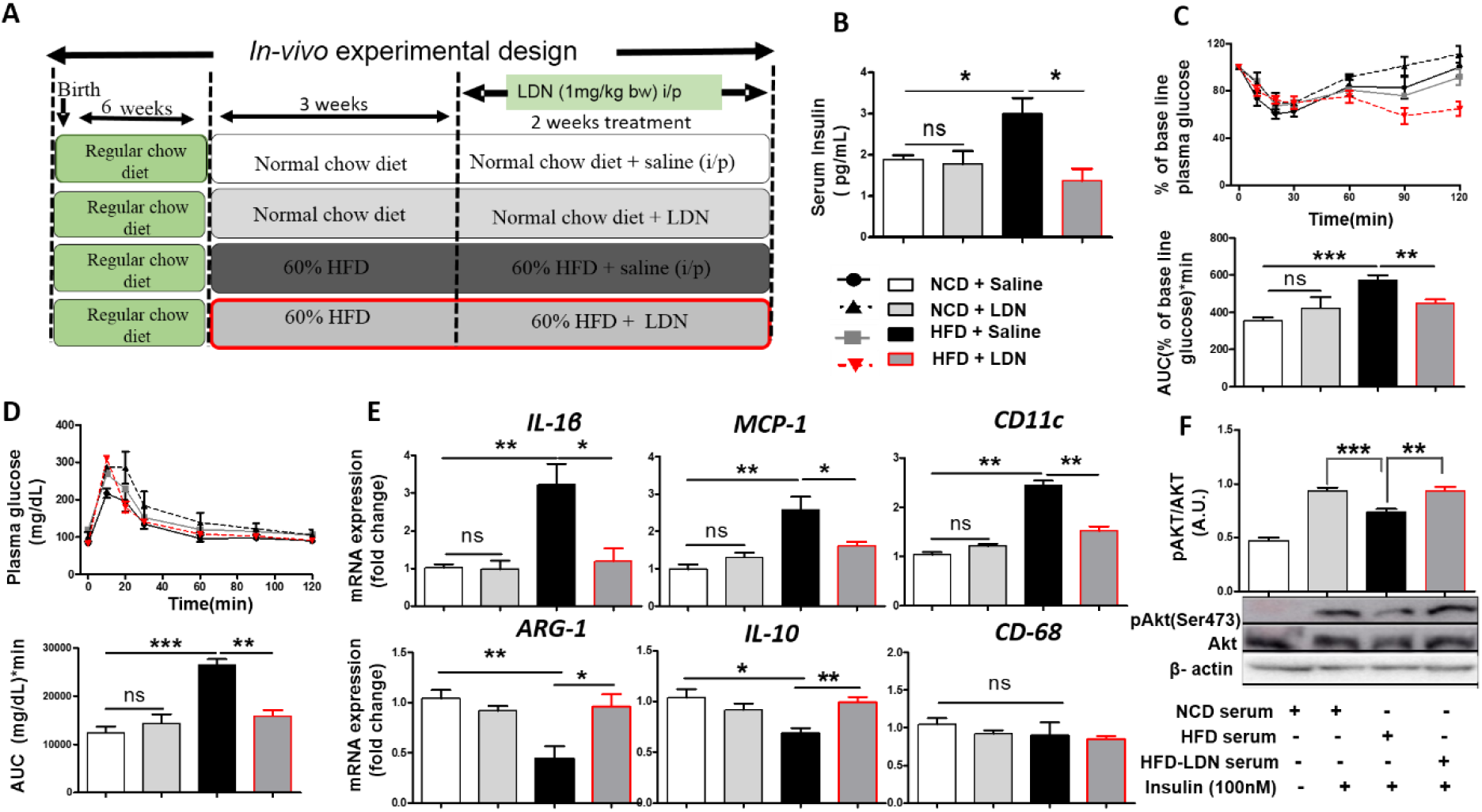
HFD induced hyperinsulinemia induces insulin resistance and secretion of M1 cykine by ATMs. **(A)** Schematic outline of short term HFD induced hyperinsulinemia mediated insulin resistance and overall *in-vivo* experimental design used in this study. (B) Fasting serum insulin levels, (C) ipITT (top) and AUC (bottom) at different time points, **(D)** Plasma Glucose level during ipGTT(top) with area under the curve (AUC) at bottom panel performed at different time in NCD-Saline, NCD-LDN, HFD-Saline, and HFD-LDN mice. **(E)** Quantitative mRNA expression of M1 (IL-1β, MCP-1, and CD11C) and M2 (ARG-1, IL-10, CD68) markers in purified ATMs from all group mice. (**F**) IB for pAKT of HepG2 cells culture in serum-free media conditioned with serum extracted from NCD-Saline, HFD-Saline, and HFD-LDN group of mice (mean ± SEM (n=3); ***p<0.001, **p<0.01 *p<0.05.)

Hence in the current study, we investigated the effect of short term HFD on cytokines expression in purified ATMs. Expression of proinflammatory markers such as *IL-1β*, *MCP-1* and *CD11c* were induced in ATMs isolated from HFD mice (Fig-2E) which was restored to normal level when mice were treated with LDN. To test our hypothesis of hyperinsulinemia promoting systemic insulin resistance by polarizing macrophage towards proinflammatory M1 phenotype and these activated macrophage release factors into systemic circulation to blunt insulin signaling in *in-vivo*, HepG2 cells were treated with serum collected from experimental mice and its effect on insulin stimulated Akt (Ser473) activation was examined. Interestingly as shown in (Fig-2F), serum from HFD-Saline fed mice blunted the Akt (Ser473) phosphorylation as compared to serum from mice fed with NCD-Saline, however, serum from HFD-LDN mice exhibited an elevated level of Akt (Ser473) phosphorylation as compared to serum from HFD-Saline mice. Naltrexone has a half-life of 4 – 10 hrs and almost fully eliminated in 24 hrs (Toljan and Vrooman, 2018), thus exclude the possibility that presence of naltrexone in serum from HFD-LDN mice interfere insulin sensitivity. These observations suggested that short term HFD treatment induced hyperinsulinemia which in turn promote ATMs to secrete M1 cytokines, and contributes to secretion of soluble factors into the systemic circulation that lead to systemic insulin resistance, whereas LDN treatment rescued from systemic insulin resistance mainly by modulating secretion of soluble factor(s) from ATM.

### Hyperinsulinemia induces release of HMGB1 in extracellular milieu

Our observations suggest that hyperinsulinemia promotes ATMs towards M1 macrophages and produce excessive quantities of proinflammatory markers in TLR4 dependent way, which in turn leads to systemic insulin resistance. However, insulin is not a ligand for TLR4, this prompted us to investigate endogenous TLR4 ligands associated with state of inflammatory dysfunction. Number of studies have demonstrated the pathophysiological connections between HMGB1 and T2DM (Wang et al., 2016). HMGB1 driven proinflammatory response have been attributed to its ability to bind and activate different cell signaling receptors such as TLR2/TLR4 (Kim et al., 2013). Thus, in current study we assessed the level of HMGB1 in hyperinsulinemic mice and found that short term HFD food regime induced release of HMGB1 to systemic circulation, whereas, LDN attenuated HFD induced serum HMGB1 level in mice (Fig-3A).

**Figure 3:**
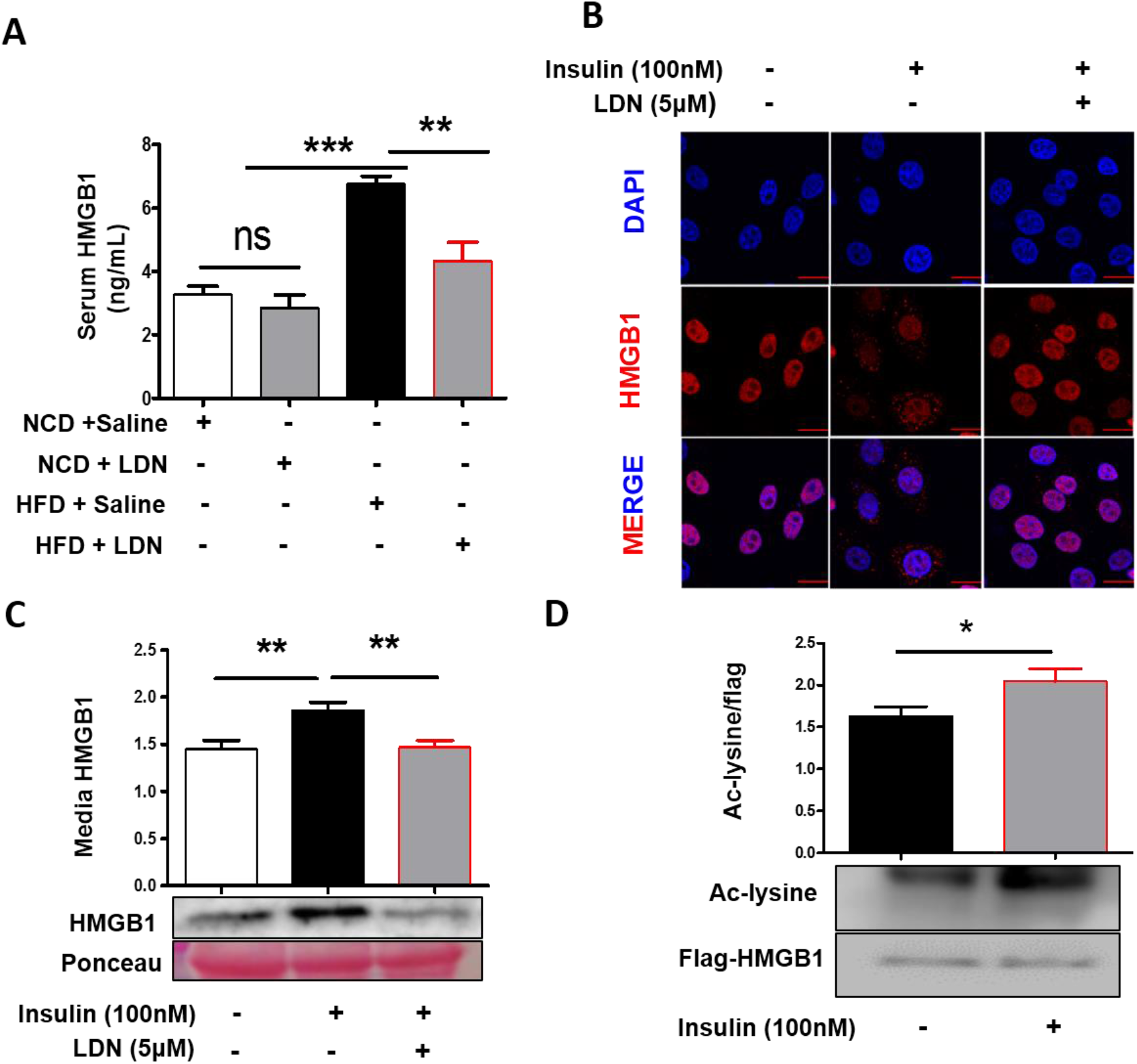
Hyperinsulinemia induced acetylation and release of HMGB1 and LDN, a TLR4 agonist significantly blocks hyperinsulinemia induced HMGB1 release. **(A)** Fasting serum HMGB1 levels in experimental mice group detected by ELISA. **(B)** immunocytochemistry for HMGB1 in HepG2 cells after 100nM insulin treatment in presence and absence of LDN for 1 hrs. showing 40X magnification, pseudocolouring: red: HMGB1,blue: nucleus counter-stain with DAPI. (**C)** HepG2 cells were incubated with LDN in presence and absence 100nM insulin and, media was collected and concentrated and blotted against anti-HMGB1. **(D)** HEK-293A cells transfected with HMGB1-Flag plasmid and allowed to grow in completed media for 48 hrs. Cells were incubated with 100nM insulin for 6hrs and cell lysate were immunoprecipitated with anti-Flag and IB with acetylated lysine and HMGB1-Flag

### Hyperinsulinemia induces Nucleus-cytosol translocation and release of HMGB1 into the extracellular milieu

HMGB1, a ubiquitous DNA-binding nuclear protein is involved in nucleosome stabilization and gene transcription (Chen et al., 2004). HMGB1 releases into the extracellular milieu during inflammation and infection. Extracellular HMGB1, plays a critical role in the development of innumerable pathological state including metabolic diseases (Wang et al., 2016). The mechanisms required for release of HMGB1 to extracellular space is translocation of HMGB1 from nucleus to cytoplasm. So, next to investigate the effect of hyperinsulinemia on HMGB1’s subcellular localization, HepG2 cells were stimulated by insulin in presence and absence of LDN. As expected, HMGB1 was predominantly expressed in the nuclei under basal condition, while we found it to be largely cytosolic in response to hyperinsulinemia (Fig-3B and S3A). Importantly nuclear localization was restored following LDN treatment (Fig-3B and S3A). Moreover, like hyperinsulinemic mice, high insulin treatment also induced HMGB1 release in the cell culture medium as examined by western blot whereas, LDN significantly blocked hyperinsulinemia-induced HMGB1 release (Fig-3C).

### Acetylation is a critical determinant of hyperinsulinemia mediated HMGB1 relocation to the cytoplasm

HMGB1 usually localized in the nucleus in basal condition and its nucleus to cytosolic translocation is a prerequisite for its release to the systemic circulation (Bonaldi, 2003; Lotze and Tracey, 2005). It is important to identify the regulation of hyperinsulinemia induced HMGB1 secretion, because HMGB1 may involve in modulating insulin signaling axis.

Post-translational modifications such as acetylation are critical for the release of HMGB1 into the extracellular milieu; as once hyperacetylated, HMGB1 gets out of nucleus and secreted out of the cell (Carneiro et al., 2009). Therefore, to determine the acetylation status of HMGB1 in hyperinsulinemic state, we transfected macrophage cell with HMGB1-flag construct and treated with insulin (100 nM) for 6 hrs and HMGB1 acetylation level in cell lysate was analyzed. Data (Fig-3D) shows an enhanced level acetylated HMGB1 in high insulin treatment. Number of reports have demonstrated that SIRT1 can directly interact and deacetylate HMGB1 to inhibit its release (Rabadi et al., 2015). We hypothesized that hyperinsulinemia mediated HMGB1 release could be modulated by SIRT1. To test this hypothesis, we measured SIRT1 protein expression profile in hyperinsulinemic state on the macrophage cells in presence and absence of LDN. Interestingly, we observed that hyperinsulinemia significantly downregulates SIRT1 protein level whereas LDN treatment protects SIRT1 from such down regulation (Fig-4A). Moreover, to dissect the role of SIRT1 on endogenous HMGB1 release, macrophage cells were treated with or without Ex-527 (SIRT1 inhibitor) (Zhao et al., 2013) for 6 hrs and HMGB1 acetylation status in media was analyzed. Data (Fig-S3B) shows that Ex-527 treatment significantly enhanced soluble acetylated HMGB1 level.

**Figure 4:**
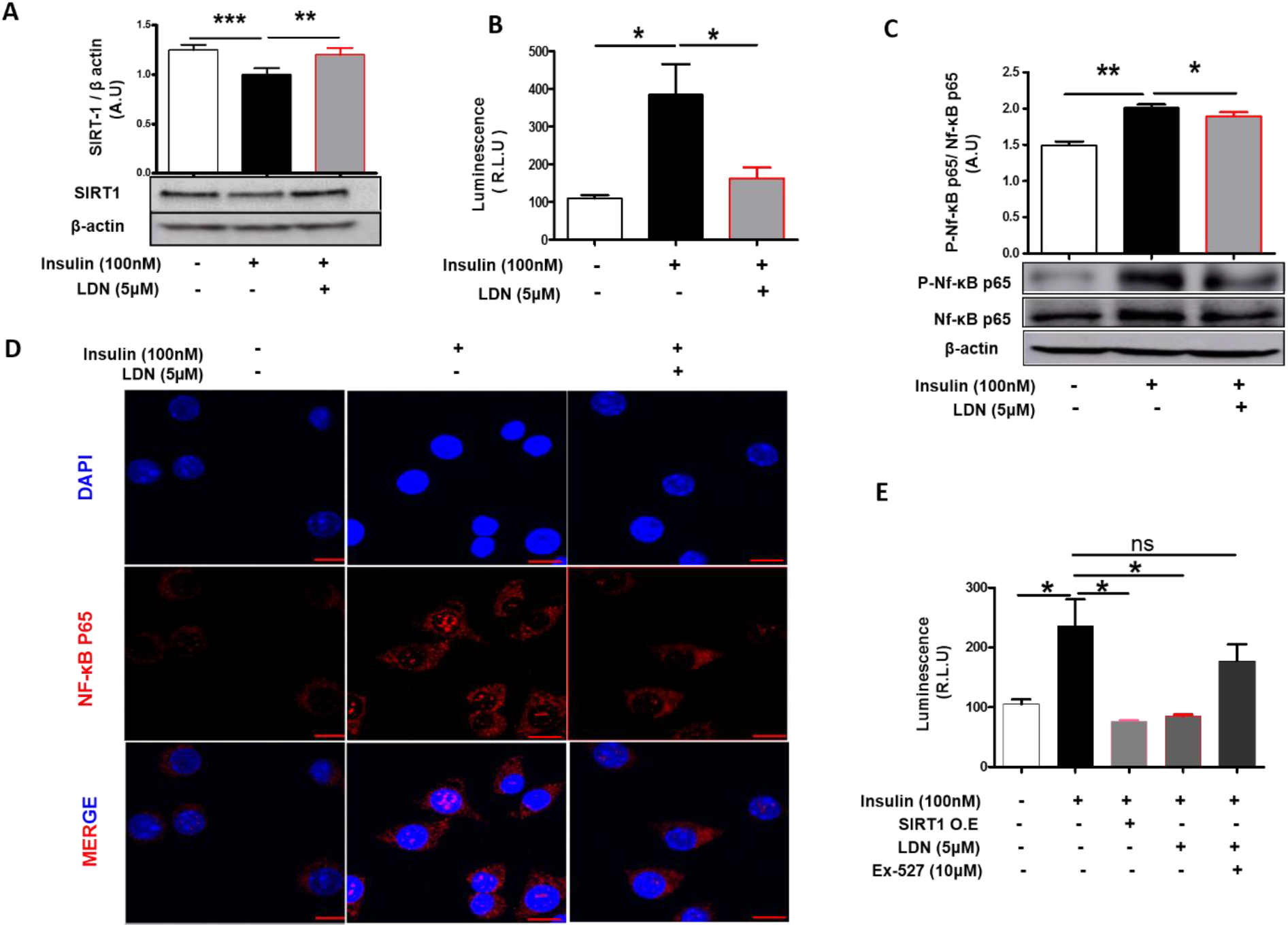
Hyperinsulinemia induced NF-κB nuclear localization in a SIRT dependent manner. **(A)** IB in Macrophage cells with high insulin in presence and absence of LDN(5μM), for 24 hours. Hyperinsulinemia downregulates SIRT1 protein whereas LDN treatment protects SIRT-1 from hyperinsulinemia mediated down regulation. **(B)** Relative light units indicating luciferase activity after transient transfection into Macrophage cells of NFκB binding site promoter-luciferase reporter constructs followed by treatment with Insulin and LDN (5μM each). Luminescence for NF-κB transcriptional activity was captured using steady glow reagent. **(C)** IB in Macrophage cells with high insulin in presence and absence of LDN for 24 hours. The cells lysate was immunoblotted against phosphor NF-κBP65. **(D)** Macrophage cells were pre-treated with or without LDN for 2 hours then stimulated with insulin (100nM) for the next 6 h. After incubation, cells were fixed and incubated against NF-κB P65 as described in methods section and imaged by confocal microscopy. **(E)** Relative light units indicating luciferase activity after transient transfection into macrophage cells of NFκB binding site promoter-luciferase reporter and SIRT1 constructs followed by treatment with Insulin and LDN, values are expressed as mean ± SEM (n=3), ***P<0.001, **P<0.01 *P<0.05.

Our study suggested that hyperinsulinemia induce downregulation of SIRT1, which in turn promotes hyperacetylation of HMGB1 and resulting in translocation of HMGB1 from nucleus to cytoplasm and LDN block hyperinsulinemia mediated HMGB1’s cytosolic localization and promotes its nuclear retention via SIRT1.

### Hyperinsulinemia induced NF-κB nuclear localization in a SIRT dependent manner

SIRT1 is also a critical factor in the negative regulation of NF-κB; a key transcription factor that drives proinflammatory phenotype, NF-κB is held quiescent in the cytoplasm in complex with IκBα (Cohen et al., 1998). In response to a proinflammatory stimulus via TLR, IκBα is phosphorylated, thereby allowing NF-κB to translocate to the nucleus and activate the transcription of a cascade of proinflammatory cytokines and chemokines to induce inflammatory responses (Yamaoka et al., 1998).

To investigate whether NF-κB transcription was regulated by hyperinsulinemia treatment, a construct containing luciferase enzyme that is driven by a NF-κB promoter was transfected to macrophage cells. We observed that hyperinsulinemia increased NF-κB transcription and LDN prevent such transactivation (Fig-4B). NF-κB activation also relies on post-translational modifications of NF-κB subunits, such as p65 phosphorylation (Viatour et al., 2005). In addition to increased transcription of NF-κB, elevated level of NF-κB p65 phosphorylation was also observed in hyperinsulinemic treatment compared to untreated cells which was attenuated in presence of LDN (Fig-4C). Phosphorylation of the p65 subunit, is essential for cytoplasm to nuclear localization of NF-κB/p65 and induce inflammatory responses. Next, we examined NF-κB/p65 localization which was barely detectable in the nucleus of unstimulated macrophage cells and as expected, hyperinsulinemia increased nuclear NF-κB localization, whereas LDN treatment blocked this shuttling of NF-κB to nucleus from cytosol (Fig-4D). Taken together, these results suggested that NF-κB was translocated into the nucleus in response to high insulin conditions *in-vitro* while, the LDN inhibit the hyperinsulinemia mediated NF-κBs’ transactivation by restricting the nuclear translocation of NF-κB. SIRT1 is known to be a positive regulator of insulin signaling and number of evidences have indicated the involvement of SIRT1 in insulin resistance and T2DM (Li et al., 2011). SIRT1 activity and/or expression is reported to be downregulated in different metabolically sensitive organs in diabetic and insulin resistant rodent model (Li et al., 2011).

To further characterize the potential role of SIRT1 in regulating LDN mediated attenuation of NF-κB activity and thereby downregulating proinflammatory response and its functional consequences, macrophage cells transfected with a NF-kB/luciferase reporter construct were treated with high insulin and LDN in presence and absence of SIRT-1 specific inhibitor (EX-527) and SIRT1 over-expression. NF-κB trans activity showed a 2.5-fold increase when cells were treated with insulin as measured by luciferase assay (Fig-4E), whereas, LDN as well SIRT1 over-expression significantly attenuated hyperinsulinemia induced NF-κB activity. Interestingly, LDN mediated suppression of NF-κB promoter activity was gone when the cells treated with EX-527 (Fig-4E). Furthermore, SIRT1 overexpression attenuated high insulin mediated proinflammatory gene expression (Fig-S4A). To further characterize the potential role of SIRT1 in regulating LDN mediated attenuation of proinflammatory gene expression, macrophage cells were treated with high insulin and LDN in presence and absence of SIRT1 specific inhibitor (EX-527) and analyzed for proinflammatory gene expression. As shown (Fig-S4B), cells co-treated with LDN and EX-527 showed no change in proinflammatory gene expression profile as compared to high insulin treated group. Taken together, these results indicate that SIRT1 positively regulates anti-inflammatory phenotype, and LDN stimulated anti-inflammatory phenotype at least partially depends on SIRT1. The SIRT1 deacetylase activity was evaluated using a deacetylase fluorometric assay kit (CS1040, Sigma-Aldrich). LDN treatment showed significantly higher SIRT1 deacetylase activity in respect to control (Fig-S4C). Together these results suggest that SIRT1 is protective against hyperinsulinemia related macrophage polarization towards M1 phenotype by silencing the transcriptional activity of NF-κB and LDN can directly enhance SIRT1 activity, and LDN mediated reduction of proinflammatory gene expression is regulated by SIRT1.

### Mechanistic connection between SIRT1 and HMGB1 translocation in genetically define mouse models

Having shown the importance of acetylation on HMGB1 for its release and SIRT1 as a critical regulator for hyperinsulinemia mediated HMGB1 release in *in-vitro*, next we aimed to evaluate if this can be experimentally proven in *in-vivo*. Towards this, we used mice model with SIRT1 overexpression in liver (Fig-5A). Physiologic endogenous insulin secretion provoked by fed state resulted in increased serum HMGB1 when compared to 24hr starved mice (Fig-5B-C). Overnight fasting and subsequently refed mice exhibited an increase in serum insulin and HMGB1 levels (Fig-5B-C). While, serum HMGB1 levels remained unchanged during overnight fasting and fed state in SIRT1 overexpressed mice model (*SIRT1OE*) (Fig-5C), indicating that the serum HMGB1 release is regulated by hyperinsulinemic cues and SIRT1 overexpression attenuates serum HMGB1 release mediated by physiologic endogenous high insulin level.

**Figure 5:**
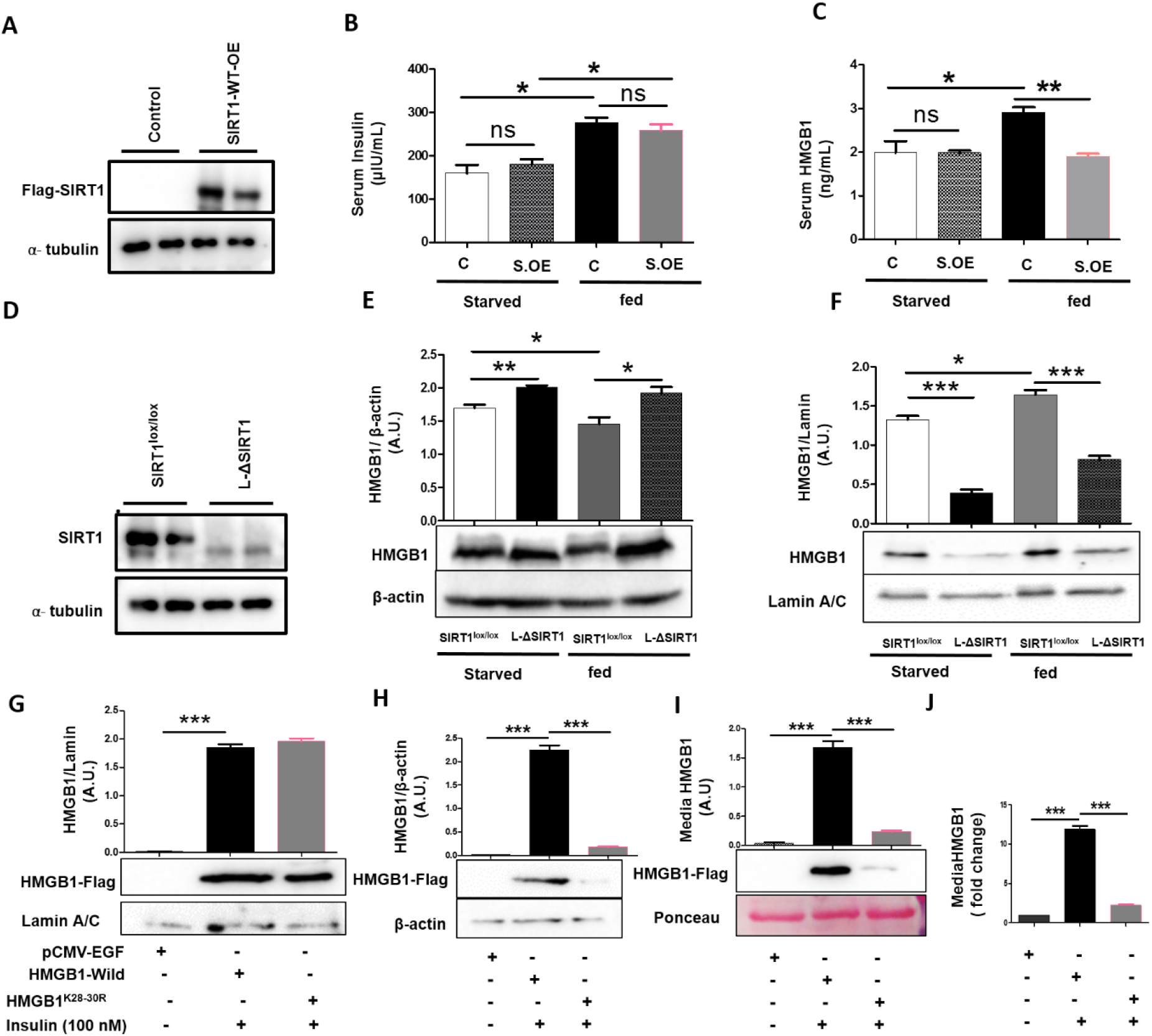
Mechanistic connection between SIRT1 and HMGB1: **(A)** Representative liver IB of control and SIRT1 over expression 18 days after adenovirus treatment. **(B)** serum insulin and **(C)** serum HMGB1 level in control and SIRT1 overexpressed mice (S. OE), serum was collected after 24h of starvation or 6h post feeding. **(D)** Representative liver IB of *Sirt1^fl/fl^* and L-ΔSIRT1 5 days after adenovirus treatment. L-ΔSIRT1 mice show SIRT1 ablation. **(E)** cytoplasmic and **(F)** Nuclear of HMGB1 protein in *Sirt1^fl/fl^* and L-ΔSIRT1 liver lysates. liver tissues were harvested either after 24h of starvation or 6h post feeding. **(G)** HEK293A cells were transfected with EGFP, HMGB1-Flag-Wild type and HMGB1^K28-30R^ –Flag mutant constructs and allowed to grow in completed media for 48 hrs. Cells were incubated with or without 100nM insulin and analysed nuclear/cytoplasmic HMGB1-Flag by fractionating cell, **(I-J)** cell supernatant were concentrated and extracellular HMGB1-Flag were determined using Western blot analysis and ELISA.

To confirm the role of the liver SIRT1 in mediating *in-vivo* effects on hepatic HMGB1 nucleo-cytoplasmic shuttling, we conditionally ablated liver SIRT1 by Adv-CRE delivery (SIRT^Δ/Δ^; L-ΔSIRT1). Adv-GFP injected *SIRT1*^lox/lox^ mice served as controls. Liver extracts from Adv-CRE injected mice revealed SIRT1 knockout (Fig-5D). To confirm that the HMGB1 protein had been translocated into the cytoplasm of L-ΔSIRT1 liver, the liver lysates from ΔSIRT1 mice were fractionated to cytoplasmic and nuclear fraction, and the distribution of HMGB1 was analyzed using western blot. L-ΔSIRT1 mice showed a robust HMGB1 cytoplasmic accumulation and minimal nuclear expression in both overnight fast and fed state compared with wild type mice (Fig-5E-5F). It is known that fasting state positively regulate SIRT1 protein and fed state downregulate SIRT1 expression (Noriega et al., 2011).

In L-ΔSIRT1 mice absence of physiologic endogenous hepatic SIRT1 induced robust HMGB1 cytoplasmic accumulation. These results reinforce the conclusion that fasting signals induce SIRT1 protein which in turn deacetylates HMGB1 at specific lysine residues and results in robust HMGB1 nuclear accumulation. Together, these findings indicate that insulin at sufficiently high concentrations induces HMGB1 nucleus to cytoplasmic shuttling and extracellular release in a SIRT1 dependent manner. Further, *in-vivo* disruption of the hepatic SIRT1 alone induces liver HMGB1 release in extracellular milieu even in absence of additional insulin signaling.

### Hyperinsulinemia-induced acetylation of HMGB1 determines its translocation from the nucleus to the cytoplasm

HMGB1 has three critical lysine residues at positions 28 to 30, known to acetylate and regulate inflammatory signaling cascades (Hwang et al., 2015). To study whether acetylation on these residue(s) is responsible for hyperinsulinemia mediated nucleo-cytosolic translocation and release of HMGB1, we generated hypo-acetylated HMGB1 mimic, where, we substituted the lysine residue with arginine at residues 28, 29, and 30 of HMGB1 (HMGB1^K28-30R^), which mimics the hypo-acetylation state (Li et al., 2007).

HEK-293A cells were transfected with wild type HMGB1-flag construct and HMGB1^K28-30R^ mutant constructs and stimulated by insulin. Hyperinsulinemia induced robust HMGB1 cytoplasmic accumulation was assessed by Western blot from wild type HMGB1, whereas, HMGB1^K28-30R^ mutant was almost undetectable in cytosolic fraction (Fig-5G-H). Moreover, cell culture medium assessed by Western blot and ELISA showed strong HMGB1 release in wild type HMGB1, however, overexpression of hypo-acetylation mimic HMGB1^K28-30R^ significantly blocked such release (Fig-5I-J).

These results suggest that lysine residues from 28, to 30 of HMGB1 are responsible for hyperinsulinemia mediated acetylation of HMGB1 and promote the nucleo-cytosolic translocation and extracellular release of HMGB1. These results suggest that SIRT1 mediated deacetylation of HMGB1 attenuates the secretion of proinflammatory cytokines, thereby may be protecting against the hyperinsulinemia induced insulin resistance.

### Neutralizing HMGB1 prevents diabetic serum to blunt insulin signaling in hepatocytes

We further examined the relevance of circulatory HMGB1 in the context of human T2DM. HMGB1 levels were increased in serum from humans with T2DM as compared to nondiabetic humans (Fig-6A). Circulating HMGB1 levels were found to be positively correlated with serum insulin level in patients with T2DM (Fig-6B). Next, we examined the relevance of serum HMGB1 in hepatic insulin signaling. Media conditioned with serum from T2DM, but not from non-diabetic individuals significantly reduced insulin-induced Akt (Ser473) phosphorylation in HepG2 cells (Fig-6C). Now to circumvent the effect of HMGB1, we neutralized mammalian diabetic serum with anti-HMGB1 antibody in equimolar concentration of HMGB1 present in serum. Interestingly, the media conditioned with HMGB1-neutrilized diabetic subject serum (ΔCM) enhanced insulin activated Akt (Ser473) phosphorylation in hepatocyte (Fig-6C). These observations indicate that increased circulating (extracellular) HMGB1 is sufficient to mitigate hepatic insulin signaling. Taken together, it suggest that hyperinsulinemia stimulates HMGB1’s translocation to cytoplasm from nucleus for its release into the systemic circulation and effectuate insulin resistance through proinflammatory responses.

**Figure 6:**
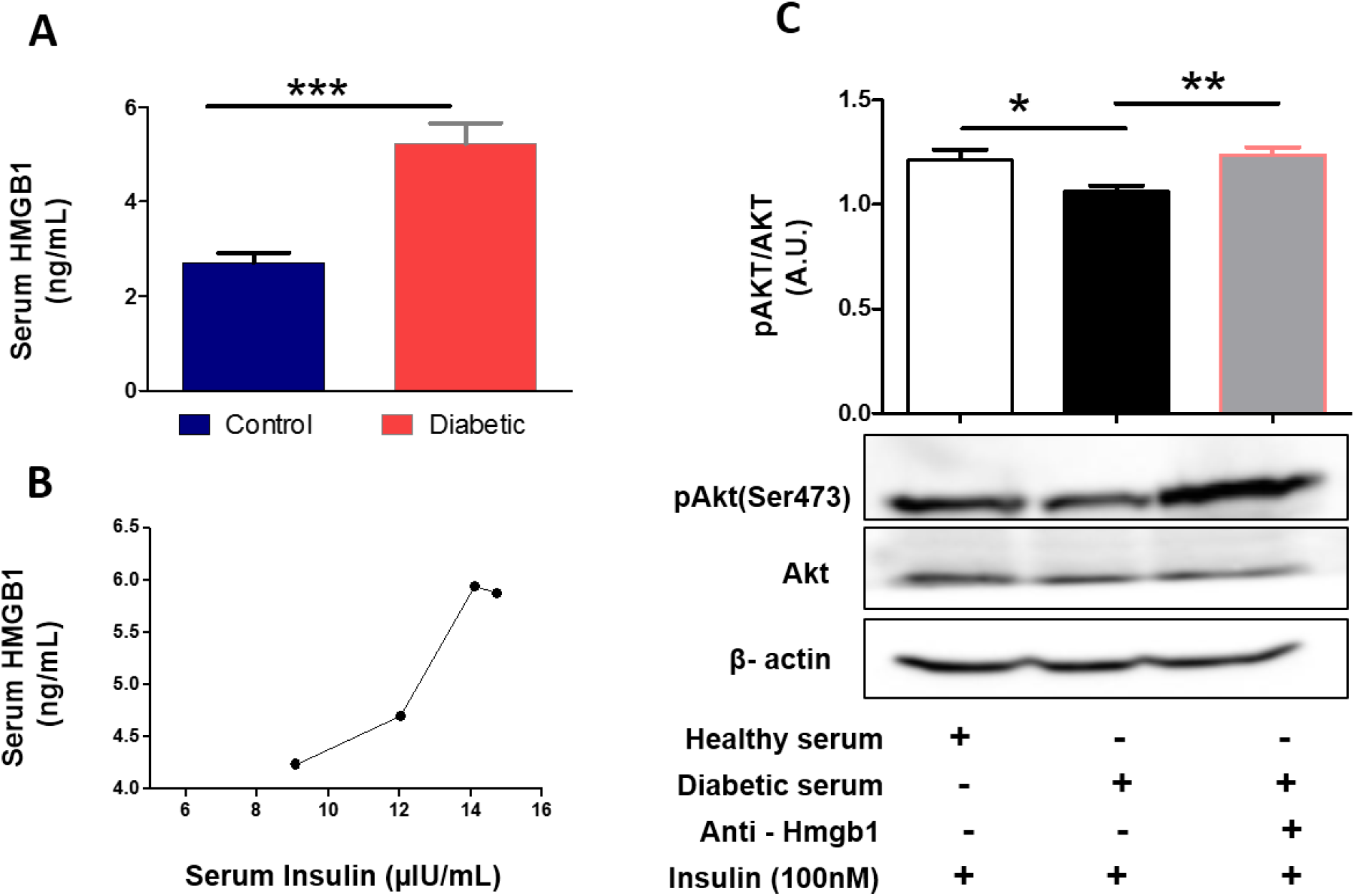
Plasma HMGB1 levels are elevated in humans with T2DM: **(A)** Serum HMGB1 level in healthy(control) and humans with T2DM, serum HMGB1 levels are elevated in humans with T2DM compared to humans without diabetes (mean±SEM; p< 0.001). **(B)** Serum HMGB1 v/s insulin in T2DM subject. **(C)** IB in HepG2 cells of HMGB1 in media conditioned with serum from healthy and T2DM subject. T2DM serum was neutralized by anti-HMGB1 antibody with equimolar concentration. IB of pAKT is suppressed during in media conditioned with T2DM serum, whereas HMGB1-neutrilized T2DM serum enhance insulin sensitivity as suggested by AKT phosphorylation (mean ±SEM; *p < 0.05), values are expressed as mean ± SEM (n=3), ***P<0.001, **P<0.01 *P<0.05.

## Discussion

Emerging data have suggested that insulin hypersecretion is not only an adaptive response to insulin resistance but may also be a primary defect (Nankervis et al., 1985; Sbraccia et al., 1996). Moreover, β-cell hypersecretion was reported as a causal agent behind the development of hepatosteatosis and insulin resistance (Steneberg et al., 2015). On the other hand, it was also reported that reduced insulin secretion can protect the mice from high fat diet mediated obesity and hyperglycemia (Mehran et al., 2012). Hypothesis of “hyperinsulinemia attenuated insulin sensitivity” was further supported by transgenic mice with extra copies of human insulin genes induced basal hyperinsulinemia and give rise to insulin resistance. however, there was no change in the body weight between transgenic and non-transgenic group (Steneberg et al., 2015(Shyng et al., 1998). In addition, *in-vitro* experiments have suggested that reducing insulin secretion from islets isolated from patients with diabetes can restore insulin pulsatility and improve pancreatic β-cell function (Laedtke et al., 2000). However, exactly how hypersecretion of insulin could lead to systemic insulin resistance and overt diabetes is not known.

Current study reported an association between hyperinsulinemia and systemic insulin resistance through promoting the release of soluble mediators from immune cells or adipose tissue, and Liver in particular. Hyperinsulinemia, which occurs early during development of T2DM, downregulates SIRT1. SIRT1 in turn modulates the extracellular release of the proinflammatory soluble mediator HMGB1 via deacetylation. HMGB1, being a TLR4 endogenous ligand, activate TLR4 dependent proinflammation which ultimately leads to insulin resistance.

Nucleocytoplasmic shuttling of HMGB1 is a significant step for its release during inflammation and numbers of post translations modification that determine its cellular localization have been elucidated (Bonaldi, 2003; Evankovich et al., 2010; Zhang et al., 2008). This study showed that. Ectopic expression of the hypo-acetylated mutant HMGB1^K28-30R^ inhibited high insulin-induced HMGB1s’ nucleo-cytoplasmic translocation and release, indicating that high insulin mediated HMGB1 nucleo-cytoplasmic translocation and release is tightly regulated by the acetylation status of these residues.

Using genetically defined mice models we also recognize liver as site of predominant sources of circulating HMGB1, and high insulin mediates increase release of HMGB1 which contributes to development of local as well as systemic insulin resistance. Our data indicates that HMGB1 release by high insulin level in the fed state, however, SIRT1 overexpression attenuates serum HMGB1 release mediated by physiologic endogenous high insulin level. The role of SIRT1 in mediating hepatic HMGB1 nucleo-cytoplasmic shuttling and release was further analysed using liver specific SIRT1 knockout mice (L-ΔSIRT1). L-ΔSIRT1 mice showed a robust HMGB1 cytoplasmic accumulation both in overnight fast and fed state compared with wild type mice. These results reinforce the conclusion that insulin at sufficiently high concentrations induces HMGB1 nucleus to cytoplasmic shuttling and extracellular release in a SIRT1 dependent manner.

Interestingly, like hypo-acetylated mutant HMGB1^K28-30R^, LDN (an opioid receptor antagonist and a known TLR4 antagonist) treatment attenuated high insulin mediated HMGB1 release. In the current study, we demonstrated that LDN inhibit HMGB1 nucleo-cytoplasmic shuttling and extracellular release. Moreover, current study also assigns an unexpected role of LDN as a SIRT1 activator, where LDN treatment markedly attenuated hyperinsulinemia induced release of HMGB1 and reduces high fat diet induced systemic insulin resistance.

Consistent with our findings, pharmacological activation of SIRT1 by resveratrol significantly reduces the extracellular release of HMGB1 (Yu et al., 2019; Zainal et al., 2017). Accordingly, LDN emerges as an activator of SIRT1 can be explored further to ameliorate the inflammation-induced diseases by decreasing the extracellular level of HMGB1.

However, uses of anti-inflammatory agents have very modest effect on improving insulin sensitivity as reported by number of studies (Larsen et al., 2007; Stanley et al., 2011). On the conjecture of current study, one can speculate that anti-inflammatory agents are failed to ameliorate the inflammation-induced insulin resistance as that these therapeutic are not fully directed at the key downstream elements, causing insulin resistance and that attempts to use LDN might be more efficacious and appear to be more rational.

Based on the studies herein, humans with T2DM exhibit increased circulating HMGB1, and their serum suppresses Akt (Ser473) phosphorylation in HepG2 cells. Our observations further suggest that neutralizing circulating HMGB1 or antagonism TLR4 by LDN enhances insulin activated Akt (Ser473) phosphorylation and insulin sensitivity. This suggests that increased circulating HMGB1 levels in T2DM may at least in part contribute to development of systemic insulin resistance to T2DM subjects.

In conclusion, our data indicates to a vicious cycle in which the hyperinsulinemia-induced repression of SIRT1 disables deacetylation of HMGB1 and facilitates its nuclear-to-cytoplasmic translocation and extracellular release thus provoke inflammation and insulin resistance which in-turn leads to hyperinsulinemia.

## ACKNOWLEDGMENTS

The authors are thankful to Director, Indian Institute of Technology Mandi for his encouragement and financial support. We sincerely thank the Advanced Materials Research Centre and BioX Centre of IIT Mandi for use of different analytical instruments and this work was partially supported by Seed Research grant from IIT Mandi to P.M. Thanks to CSIR-IITR, TIFR Mumbai, PGIMER for a research facility and AC and SD thank to Department of Science & Technology (DST) INSPIRE fellowship, KG thanks the UGC, India for her research fellowship.

## AUTHOR CONTRIBUTIONS

AC conducted majority of the experiments and analysed the data.. AC, SK, AKK, TC conducted mice experiments. KG, SD performed few in vitro experiments.BM and AB facilities clinical studies. P.M, designed study and supervised the project. PM, AC, wrote the manuscript with support from UKS, CM, and D.G and All authors discussed the results and commented on the manuscript. P.M is the guarantor of this work and, as such, has full access to all the data in the study and takes responsibility for the integrity of the data and the accuracy of the data analysis.

## Methods

### Drugs and Chemicals

A pharmacological low dose of naltrexone hydrochloride (LDN) obtained from MP Biomedicals (151725) was used.

### Cell lines and culture treatment

Human hepatocellular carcinoma (HepG2), HEK293A and murine macrophage (Raw264.7) cell lines were cultured in DMEM and RPMI media supplemented with 10% fetal bovine serum and 1% penicillin-streptomycin and MEM non-essential amino acids solution (wherever required).

### Cell viability

**C**ell viability assay was carried out in both HepG2 and Raw264.7 cells using MTT dye (3-(4, 5-dimethyl thiazol-2yl)-2, 5-diphenyl tetrazolium bromide). Cells were seeded in 96-well plate and allowed to grow overnight. Further cells were treated with varying doses (0, 2, 4, 6, 8, 10, 20, 40 μM) of the LDN for 24 hrs. Post-treatment, 10μl of MTT (5mg/ml stock in PBS) was added to all the wells. The formazan crystals thus formed were solubilized in 200μl DMSO and the absorbance was recorded using (Infinite M200 Pro TECAN).

### RNA isolation and gene expression profile

Raw264.7 cultured cells were treated with 5μM of LDN in presence and absence of 100nM insulin for 24 hrs. RNA was isolated from the cells using RNA-Xpress reagent (HIMEDIA-MB601) and 1μg of RNA was reverse-transcribed for quantitative RT-PCR analysis. The results were normalized using housekeeping genes 18S (rRNA) and expression was calculated as fold change by normalizing the control values. Primers are provided in supplemental experimental procedures.

### Western Blot Analysis

Raw264.7 cultured cells were incubated with LDN (0 and 5μM) in presence or absence of insulin (100nM) for 24 hrs and protein expression of various insulin receptor downstream such as SIRT1, NF-κB was studied. Furthermore, the phosphorylation level of Akt (Ser473), was evaluated by treating conditioned media (supernatant from insulin challenged macrophage cells), and serum (from all experimental group of mice and clinical samples) to HepG2 cells for 12 hrs. After treatment, cells were lysed in RIPA buffer containing 1% protease-phosphatase inhibitors. Protein concentration was determined by BCA assay reagent as described in manufacturer’s (Thermo Scientific-23227) manual. Protein was loaded on SDS-PAGE and electro-blotted on to PVDF membranes. The membrane was incubated in 5% milk blocking solution for 2 hrs at room temperature (RT) and probed against primary antibody (1:2000 diluted in TBST; After washing with TBST, membrane was incubated with HRP conjugated IgG secondary antibody for 2 hrs and visualized by chemiluminescence.

### NF κB reporter assay

Macrophage cells seeded in 24 well plate were transfected with a NFκB specific firefly luciferase reporter plasmid, a negative control containing non-inducible firefly luciferase reporter and a positive control containing a constitutively expressing GFP firefly luciferase construct with or without SIRT1 plasmid using lipofectamine p3000 transfection reagent (Invitrogen-L3000-015). Further, the cells were scrapped and seeded in white bottom 96 well plate, allowed to adhere properly overnight and treated with or without 100 nM of insulin in presence and absence of LDN and EX-527 for 24 hrs. The resulting cells were subsequently used for NFκB activity measurement by adding steady Glo reagent and measuring the luminescence after 5 min of incubation at RT.

### Immunocytochemistry

After treatment, cells were washed with PBS and fixed with formalin at RT. Further cells were washed with PBST (0.05% Tween 20 in PBS) to remove traces formalin. Blocking buffer (2% FBS in PBST) was added and allowed to shake moderately for 2 hrs at RT. Incubated the cells with primary antibody (1:500 dilution) overnight at 4°C. After washing with PBST (0.1% Tween-20), 2 hrs incubation with Alexa-Fluor Secondary antibodies at 1:1000 dilution was done. Slide containing cells were washed for 3 times with PBST and mounted in DAPI mounting media (Vectorshield-H1200). Image was recorded by using confocal (Nikon) microscopy.

### Animals and treatments

C57BL/6 mice (Male, six-week-old) procured from the animal house of Indian Institute of Toxicology Research (IITR), Lucknow, India and were kept as ad libitum on a normal chow diet(NCD) and 60% high-fat diet(HFD) for 3 weeks. Mice were randomly divided into 4 groups. <u>Group I (NCD):</u> Mice receiving NCD and water ad libitum up to 3 weeks and subjected to normal saline (i.p) on 4^th^ and 5^th^ week. <u>Group II (NCD - LDN):</u> Mice receiving NCD and water ad libitum up to 3 weeks and subjected to LDN (1mg/kgbw. i.p) treatment on 4^th^ and 5^th^ week. <u>Group III (HFD-Saline):</u> Mice receiving HFD and water ad libitum up to 3 weeks and subjected to normal saline (i.p) on 4^th^ and 5 ^th^ week. <u>Group IV (HFD-LDN):</u> Mice receiving HFD and water ad libitum up to 3 weeks and subjected to LDN (1mg/kgbw. i.p) on 4^th^ and 5th week. LDN treatment (1mg/kg bw) was started on 4^th^week for 14 days. Metabolic parameters were done on 11^th^ and 13^th^ day of treatment respectively.

### Genetically defined SIRT1 mice models

C57BL/6 and SIRT1^lox/lox^ mice were housed under standard animal house conditions. male mice were used for experiments. The procedures and the project were approved and were in accordance with the institute animal ethics committee (IAEC) guidelines. Mice were given injections intravenous (i.v.), via the tail vein, of Ad-SIRT1-WT or AdCMV in C57BL/6 mice, on Day-1. Liver tissues were harvested on Day-18, 6 hrs post-feeding for gene expression and western blot analysis. For SIRT1^lox/lox^ mice, adenovirus overexpressing Cre-recombinase was injected i.v. on Day 1, followed by Ad-SIRT1-WT or Ad-CMV administration on Day 5. Post injection, experiments were performed and liver tissues were harvested either after 24h of starvation or 6 hrs post feeding (6h-RF). Dissected tissues were either snap frozen in liquid nitrogen or processed for protein lysates and total RNA extraction.

### Tissue lysate preparation

Tissue lysates were prepared in RIPA buffer (50mM Tris pH 8.0, 150mM NaCl, 0.1% SDS, 0.5% Sodium deoxycholate, 1% Triton X 100, 0.1% SDS, 1 mM PMSF, Protease inhibitor cocktail) and debris was pelleted at 12,000 rpm (4°C/10-minutes), followed by determining protein concentration using BCA assay kit (SIGMA). Samples were loaded onto SDS-PAGE for protein expression analysis.

### Metabolic Parameters Analysis

Intraperitoneal glucose tolerance tests (ipGTT) and insulin tolerance tests (ipITT) were performed according to standard protocols(Mondal et al., 2015). Animals were administered 20% D-glucose (2 g/kg orally via gavage) or insulin (0.5 units/kg ip.) recombinant human insulin, respectively. Tail-vein blood was collected at the indicated times in figures for glucose and insulin measurements. serum glucose was measured using a glucometer. Serum insulin and HMGB1 level were measured using insulin and HMGB1 ELISA.

### ATM Isolation and Analysis

Mice were euthanized chemically and epididymal fat was processed for isolation of ATM. 1 gram of adipose fat was rinsed in PBS and minced to small pieces in HEPES-DMEM buffer containing 10 mg/ml BSA. The suspension was centrifuged at 1,000g for 10 min and the resultant supernatant was pipette off to fresh tubes. 1mg/ml of collagenase type-IV and 50U/ml DNAse-II were added to this suspension and incubated at 37°C for 45 mins with moderate shaking, filtered through 250-micron filter and the resultant solution was centrifuged again at 1,000g for 10 mins. Floating cells contained adipocyte and pellet are SVC. RBC lysis buffer was added gently to disrupt the sedimented pellets and centrifuged at 1,000 g for 10 min at 4°C. Fat macrophage cells were isolated by using BD IMag anti-mouse CD11b+ magnetic beads through positive selection under magnetic field. The percentage purity of macrophage isolation was determined by FACSCANTO II flow cytometer using APC tagged CD11b monoclonal antibody. The isolated macrophage was processed for RNA isolation, cDNA synthesis and various M1-M2 markers were evaluated using real time PCR.

### Subcellular Fractionation

Cultured cells and liver tissue were separated into cytoplasmic and nuclear fractions using NE-PER (Thermo Scientific-78833) and all the centrifugation steps were carried at 4°C. Immunoblot analysis was performed in cytoplasmic and nuclear fraction using respective antibodies.

### HMGB1 Neutralization

Mammalian diabetic serum was neutralized by anti-HMGB1 antibody in equimolar concentration of hmgb1 present in serum and challenged to HepG2 cells. Further the phosphorylation of Akt (Ser473) was assessed.

### Clinical experimentation and subject details

The experimental protocol was reviewed and approved from Institutional Ethics Committee, PGIMER-Chandigarh. All the participants/subjects were screened and enrolled as per the inclusion/exclusion criteria (for detail see characteristics of subjects in supplementary information). Informed consent was obtained from all the subjects enrolled (including healthy volunteers) and provided with instructions for 1. Overnight fasting, 2. No physical activity. 3. No other exertion in order to avoid stress related factor. All the participants were analyzed for several metabolic parameters such as fasting glucose level (FG), fasting basal insulin (FI), fasting HbA1c, fasting C-peptide) in the respective department of PGIMER-Chandigarh. Blood samples were collected in BD Vacutainer^®^ plastic tube, centrifuged to get serum and stored in aliquots at −80°C till further use.

### Statistical analysis

All the data presented is as mean ± SEM of three individual experiments unless specified. Comparisons between means were performed using Student *t*-test for unpaired data within two conditions.

## References

Badman, M.K., Pissios, P., Kennedy, A.R., Koukos, G., Flier, J.S., and Maratos-Flier, E. (2007). Hepatic Fibroblast Growth Factor 21 Is Regulated by PPARα and Is a Key Mediator of Hepatic Lipid Metabolism in Ketotic States. Cell Metab. 5, 426–437.

Blüher, M. (2012). Clinical Relevance of Adipokines. Diabetes Metab. J. 36, 317.

Blüher, M., and Mantzoros, C.S. (2015). From leptin to other adipokines in health and disease: Facts and expectations at the beginning of the 21st century. Metabolism 64, 131–145.

Bonaldi, T. (2003). Monocytic cells hyperacetylate chromatin protein HMGB1 to redirect it towards secretion. EMBO J. 22, 5551–5560.

Carneiro, V.C., de Moraes Maciel, R., de Abreu da Silva, I.C., da Costa, R.F.M., Paiva, C.N., Bozza, M.T., and Fantappié, M.R. (2009). The extracellular release of Schistosoma mansoni HMGB1 nuclear protein is mediated by acetylation. Biochem. Biophys. Res. Commun. 390, 1245–1249.

Chen, G., Ward, M.F., Sama, A.E., and Wang, H. (2004). Extracellular HMGB1 as a Proinflammatory Cytokine. J. Interf. Cytokine Res. 24, 329–333.

Chen, Y.-H., Lee, Y.-C., Tsao, Y.-C., Lu, M.-C., Chuang, H.-H., Yeh, W.-C., Tzeng, I.-S., and Chen, J.-Y. (2018). Association between high-fasting insulin levels and metabolic syndrome in non-diabetic middle-aged and elderly populations: a community-based study in Taiwan. BMJ Open 8, e016554.

Cohen, L., Henzel, W.J., and Baeuerle, P.A. (1998). IKAP is a scaffold protein of the IκB kinase complex. Nature 395, 292–296.

Dogra, S., Kar, A.K., Girdhar, K., Daniel, P.V., Chatterjee, S., Choubey, A., Ghosh, S., Patnaik, S., Ghosh, D., and Mondal, P. (2019). Zinc oxide nanoparticles attenuate hepatic steatosis development in high-fat-diet fed mice through activated AMPK signaling axis. Nanomedicine Nanotechnology, Biol. Med. 17, 210–222.

Evankovich, J., Cho, S.W., Zhang, R., Cardinal, J., Dhupar, R., Zhang, L., Klune, J.R., Zlotnicki, J., Billiar, T., and Tsung, A. (2010). High Mobility Group Box 1 Release from Hepatocytes during Ischemia and Reperfusion Injury Is Mediated by Decreased Histone Deacetylase Activity. J. Biol. Chem. 285, 39888–39897.

Garvey, W.T., Huecksteadt, T.P., Matthaei, S., and Olefsky, J.M. (1988). Role of glucose transporters in the cellular insulin resistance of type II non-insulin-dependent diabetes mellitus. J. Clin. Invest. 81, 1528–1536.

Hwang, J.S., Choi, H.S., Ham, S.A., Yoo, T., Lee, W.J., Paek, K.S., and Seo, H.G. (2015). Deacetylation-mediated interaction of SIRT1-HMGB1 improves survival in a mouse model of endotoxemia. Sci. Rep. 5, 15971.

Kim, S., Kim, S.Y., Pribis, J.P., Lotze, M., Mollen, K.P., Shapiro, R., Loughran, P., Scott, M.J., and Billiar, T.R. (2013). Signaling of High Mobility Group Box 1 (HMGB1) through Toll-like Receptor 4 in Macrophages Requires CD14. Mol. Med. 19, 88–98.

Klune, J.R., Dhupar, R., Cardinal, J., Billiar, T.R., and Tsung, A. (2008). HMGB1: Endogenous Danger Signaling. Mol. Med. 14, 476–484.

Laedtke, T., Kjems, L., Pørksen, N., Schmitz, O., Veldhuis, J., Kao, P.C., and Butler, P.C. (2000). Overnight inhibition of insulin secretion restores pulsatility and proinsulin/insulin ratio in type 2 diabetes. Am. J. Physiol. Metab. 279, E520–E528.

Larsen, C.M., Faulenbach, M., Vaag, A., Vølund, A., Ehses, J.A., Seifert, B., Mandrup-Poulsen, T., and Donath, M.Y. (2007). Interleukin-1–Receptor Antagonist in Type 2 Diabetes Mellitus. N. Engl. J. Med. 356, 1517–1526.

Li, X., Zhang, S., Blander, G., Tse, J.G., Krieger, M., and Guarente, L. (2007). SIRT1 deacetylates and positively regulates the nuclear receptor LXR. Mol. Cell 28, 91–106.

Li, Y., Xu, S., Giles, A., Nakamura, K., Lee, J.W., Hou, X., Donmez, G., Li, J., Luo, Z., Walsh, K., et al. (2011). Hepatic overexpression of SIRT1 in mice attenuates endoplasmic reticulum stress and insulin resistance in the liver. FASEB J. 25, 1664–1679.

Lotze, M.T., and Tracey, K.J. (2005). High-mobility group box 1 protein (HMGB1): nuclear weapon in the immune arsenal. Nat. Rev. Immunol. 5, 331–342.

Marín-Juez, R., Jong-Raadsen, S., Yang, S., and Spaink, H.P. (2014). Hyperinsulinemia induces insulin resistance and immune suppression via Ptpn6/Shp1 in zebrafish. J. Endocrinol. 222, 229–241.

Mehran, A.E., Templeman, N.M., Brigidi, G.S., Lim, G.E., Chu, K.-Y., Hu, X., Botezelli, J.D., Asadi, A., Hoffman, B.G., Kieffer, T.J., et al. (2012). Hyperinsulinemia Drives Diet-Induced Obesity Independently of Brain Insulin Production. Cell Metab. 16, 723–737.

Misu, H., Takamura, T., Takayama, H., Hayashi, H., Matsuzawa-Nagata, N., Kurita, S., Ishikura, K., Ando, H., Takeshita, Y., Ota, T., et al. (2010). A Liver-Derived Secretory Protein, Selenoprotein P, Causes Insulin Resistance. Cell Metab. 12, 483–495.

Mondal, P., Song, W.-J., Li, Y., Yang, K.S., and Hussain, M.A. (2015). Increasing β-cell mass requires additional stimulation for adaptation to secretory demand. Mol. Endocrinol. 29, 108–120.

Nankervis, A., Proietto, J., Aitken, P., and Alford, F. (1985). Hyperinsulinaemia and insulin insensitivity: studies in subjects with insulinoma. Diabetologia 28, 427–431.

Noriega, L.G., Feige, J.N., Canto, C., Yamamoto, H., Yu, J., Herman, M.A., Mataki, C., Kahn, B.B., and Auwerx, J. (2011). CREB and ChREBP oppositely regulate SIRT1 expression in response to energy availability. EMBO Rep. 12, 1069–1076.

Olefsky, J.M., Garvey, W.T., Henry, R.R., Brillon, D., Matthael, S., and Freidenberg, G.R. (1988). Cellular mechanisms of insulin resistance in non-insulin-dependent (type II) diabetes. Am. J. Med. 85, 86–105.

Pal, D., Dasgupta, S., Kundu, R., Maitra, S., Das, G., Mukhopadhyay, S., Ray, S., Majumdar, S.S., and Bhattacharya, S. (2012). Fetuin-A acts as an endogenous ligand of TLR4 to promote lipid-induced insulin resistance. Nat. Med. 18, 1279–1285.

Pereira, S., Park, E., Mori, Y., Haber, C.A., Han, P., Uchida, T., Stavar, L., Oprescu, A.I., Koulajian, K., Ivovic, A., et al. (2014). FFA-induced hepatic insulin resistance in vivo is mediated by PKCδ, NADPH oxidase, and oxidative stress. Am. J. Physiol. Metab. 307, E34–E46.

Pfluger, P.T., Herranz, D., Velasco-Miguel, S., Serrano, M., and Tschop, M.H. (2008). Sirt1 protects against high-fat diet-induced metabolic damage. Proc. Natl. Acad. Sci. 105, 9793–9798.

Rabadi, M.M., Xavier, S., Vasko, R., Kaur, K., Goligorksy, M.S., and Ratliff, B.B. (2015). High-mobility group box 1 is a novel deacetylation target of Sirtuin1. Kidney Int. 87, 95–108.

Reaven, G.M. (2005). THE INSULIN RESISTANCE SYNDROME: Definition and Dietary Approaches to Treatment. Annu. Rev. Nutr. 25, 391–406.

Rhee, E.-J., Lee, W.-Y., Cho, Y.-K., Kim, B.-I., and Sung, K.-C. (2011). Hyperinsulinemia and the Development of Nonalcoholic Fatty Liver Disease in Nondiabetic Adults. Am. J. Med. 124, 69–76.

Rickenbacher, A., Jang, J.H., Limani, P., Ungethüm, U., Lehmann, K., Oberkofler, C.E., Weber, A., Graf, R., Humar, B., and Clavien, P.-A. (2014). Fasting protects liver from ischemic injury through Sirt1-mediated downregulation of circulating HMGB1 in mice. J. Hepatol. 61, 301–308.

Sbraccia, P., D’Adamo, M., Leonetti, F., Caiola, S., Iozzo, P., Giaccari, A., Buongiorno, A., and Tamburrano, G. (1996). Chronic primary hyperinsulinaemia is associated with altered insulin receptor mRNA splicing in muscle of patients with insulinoma. Diabetologia 39, 220–225.

Shin, K.C., Hwang, I., Choe, S.S., Park, J., Ji, Y., Kim, J.I., Lee, G.Y., Choi, S.H., Ching, J., Kovalik, J.-P., et al. (2017). Macrophage VLDLR mediates obesity-induced insulin resistance with adipose tissue inflammation. Nat. Commun. 8, 1087.

Shoelson, S.E., Lee, J., and Goldfine, A.B. (2006). Inflammation and insulin resistance. J. Clin. Invest. 116, 1793–1801.

Shyng, S.L., Ferrigni, T., Shepard, J.B., Nestorowicz, A., Glaser, B., Permutt, M.A., and Nichols, C.G. (1998). Functional analyses of novel mutations in the sulfonylurea receptor 1 associated with persistent hyperinsulinemic hypoglycemia of infancy. Diabetes 47, 1145–1151.

Stanley, T.L., Zanni, M. V., Johnsen, S., Rasheed, S., Makimura, H., Lee, H., Khor, V.K., Ahima, R.S., and Grinspoon, S.K. (2011). TNF-α Antagonism with Etanercept Decreases Glucose and Increases the Proportion of High Molecular Weight Adiponectin in Obese Subjects with Features of the Metabolic Syndrome. J. Clin. Endocrinol. Metab. 96, E146–E150.

Stefan, N., and Häring, H.-U. (2013). The role of hepatokines in metabolism. Nat. Rev. Endocrinol. 9, 144–152.

Steneberg, P., Sykaras, A.G., Backlund, F., Straseviciene, J., Söderström, I., and Edlund, H. (2015). Hyperinsulinemia enhances hepatic expression of the fatty acid transporter Cd36 and provokes hepatosteatosis and hepatic insulin resistance. J. Biol. Chem. 290, 19034–19043.

Su, Z., Zhang, P., Yu, Y., Lu, H., Liu, Y., Ni, P., Su, X., Wang, D., Liu, Y., Wang, J., et al. (2016). HMGB1 Facilitated Macrophage Reprogramming towards a Proinflammatory M1-like Phenotype in Experimental Autoimmune Myocarditis Development. Sci. Rep. 6, 21884.

Sung, K.-C.C., Seo, M.-H.H., Rhee, E.-J.J., and Wilson, A.M. (2011). Elevated fasting insulin predicts the future incidence of metabolic syndrome: a 5-year follow-up study. Cardiovasc. Diabetol. 10, 108.

Thomas, J.O. (2001). HMG1 and 2: architectural DNA-binding proteins. Biochem. Soc. Trans. 29, 395–401.

Toljan, K., and Vrooman, B. (2018). Low-Dose Naltrexone (LDN)—Review of Therapeutic Utilization. Med. Sci. 6, 82.

Viatour, P., Merville, M.-P., Bours, V., and Chariot, A. (2005). Phosphorylation of NF-κB and IκB proteins: implications in cancer and inflammation. Trends Biochem. Sci. 30, 43–52.

Wang, Y., Zhong, J., Zhang, X., Liu, Z., Yang, Y., Gong, Q., and Ren, B. (2016). The Role of HMGB1 in the Pathogenesis of Type 2 Diabetes. J. Diabetes Res. 2016, 1–11.

Weerts, E.M., Kim, Y.K., Wand, G.S., Dannals, R.F., Lee, J.S., Frost, J.J., and McCaul, M.E. (2008). Differences in δ- and μ-Opioid Receptor Blockade Measured by Positron Emission Tomography in Naltrexone-Treated Recently Abstinent Alcohol-Dependent Subjects. Neuropsychopharmacology 33, 653–665.

Weisberg, S.P., McCann, D., Desai, M., Rosenbaum, M., Leibel, R.L., and Ferrante, A.W. (2003). Obesity is associated with macrophage accumulation in adipose tissue. J. Clin. Invest. 112, 1796–1808.

Xu, H., Barnes, G.T., Yang, Q., Tan, G., Yang, D., Chou, C.J., Sole, J., Nichols, A., Ross, J.S., Tartaglia, L.A., et al. (2003). Chronic inflammation in fat plays a crucial role in the development of obesity-related insulin resistance. J. Clin. Invest. 112, 1821–1830.

Yamaoka, S., Courtois, G., Bessia, C., Whiteside, S.T., Weil, R., Agou, F., Kirk, H.E., Kay, R.J., and Israël, A. (1998). Complementation cloning of NEMO, a component of the IkappaB kinase complex essential for NF-kappaB activation. Cell 93, 1231–1240.

Yoshizaki, T., Milne, J.C., Imamura, T., Schenk, S., Sonoda, N., Babendure, J.L., Lu, J.-C., Smith, J.J., Jirousek, M.R., and Olefsky, J.M. (2009). SIRT1 Exerts Anti-Inflammatory Effects and Improves Insulin Sensitivity in Adipocytes. Mol. Cell. Biol. 29, 1363–1374.

Yu, S., Zhou, X., Xiang, H., Wang, S., Cui, Z., and Zhou, J. (2019). Resveratrol Reduced Liver Damage After Liver Resection in a Rat Model by Upregulating Sirtuin 1 (SIRT1) and Inhibiting the Acetylation of High Mobility Group Box 1 (HMGB1). Med. Sci. Monit. 25, 3212–3220.

Zainal, N., Chang, C.-P., Cheng, Y.-L., Wu, Y.-W., Anderson, R., Wan, S.-W., Chen, C.-L., Ho, T.-S., AbuBakar, S., and Lin, Y.-S. (2017). Resveratrol treatment reveals a novel role for HMGB1 in regulation of the type 1 interferon response in dengue virus infection. Sci. Rep. 7, 42998.

Zhang, X., Wheeler, D., Tang, Y., Guo, L., Shapiro, R.A., Ribar, T.J., Means, A.R., Billiar, T.R., Angus, D.C., and Rosengart, M.R. (2008). Calcium/Calmodulin-Dependent Protein Kinase (CaMK) IV Mediates Nucleocytoplasmic Shuttling and Release of HMGB1 during Lipopolysaccharide Stimulation of Macrophages. J. Immunol. 181, 5015–5023.

Zhao, X., Allison, D., Condon, B., Zhang, F., Gheyi, T., Zhang, A., Ashok, S., Russell, M., MacEwan, I., Qian, Y., et al. (2013). The 2.5 Å Crystal Structure of the SIRT1 Catalytic Domain Bound to Nicotinamide Adenine Dinucleotide (NAD ^+^) and an Indole (EX527 Analogue) Reveals a Novel Mechanism of Histone Deacetylase Inhibition. J. Med. Chem. 56, 963–969.

