## Supplemental document for "Hyperinsulinemia promotes HMGB1 release leading to inflammation induced systemic insulin resistance: An interplay between pancreatic beta-cell and peripheral organs"

**SiRNA transfection:**

Macrophage cells seeded in a 6 well plate were transfected with 20nM negative control siRNA and TLR4-siRNA (Ambion- 4390771) using lipofectamine RNAiMAX (Invitrogen-13778-075) and allowed to grow for 30hrs in complete media. Cells were then treated with or without insulin (100nM) for 24hrs and mRNA expression of different macrophage M1 markers were investigated using syber green based real-time PCR.

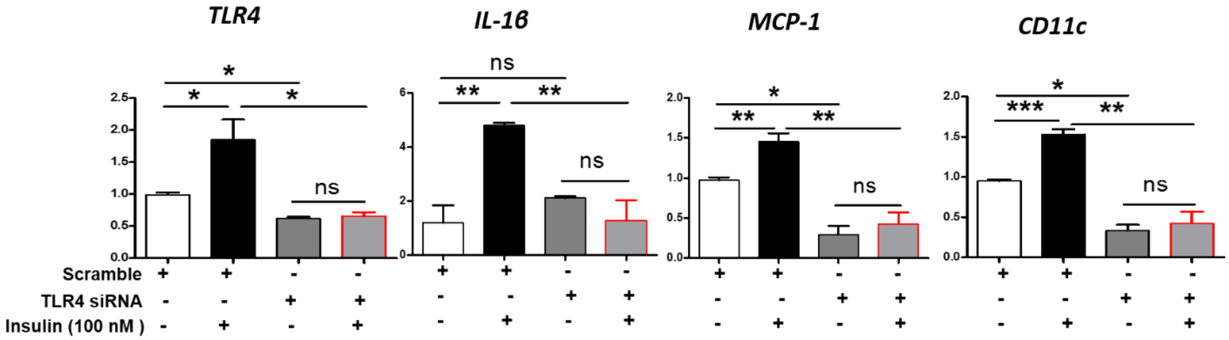


**Figure S1: Hyperinsulinemia induces proinflammatory genes expression requires TLR4:**

Quantitative RT-PCR of indicated genes ( *TLR4, IL-1β, MCP-1, CD11c*) in TLR4 knockdown murine macrophage. Cells treated with high insulin showed induced expression of proinflammatory genes. However, in cells transfected with TLR4-siRNA, high insulin mediated proinflammatory genes expression was abrogated.

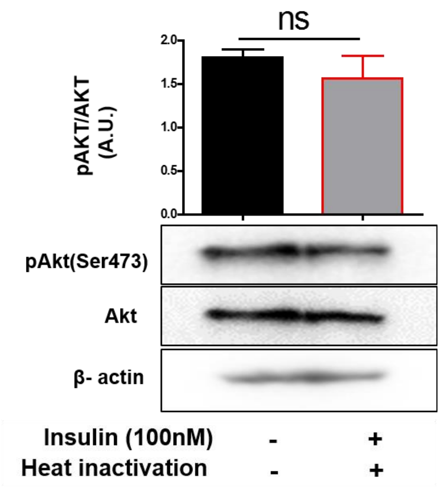


**Figure S2: Heat inactivation of hyperinsulinemia challenged macrophage conditioned media did not blunt Akt (Ser473) phosphorylation in HepG2 cells:** IB in HepG2 Cell lysates using anti pAkt (ser473), Akt and β-actin antibodies after hyperinsulinemia challenged macrophage conditioned media treatment (with or without heat inactivation), Values are expressed as mean ± SEM (n=3)

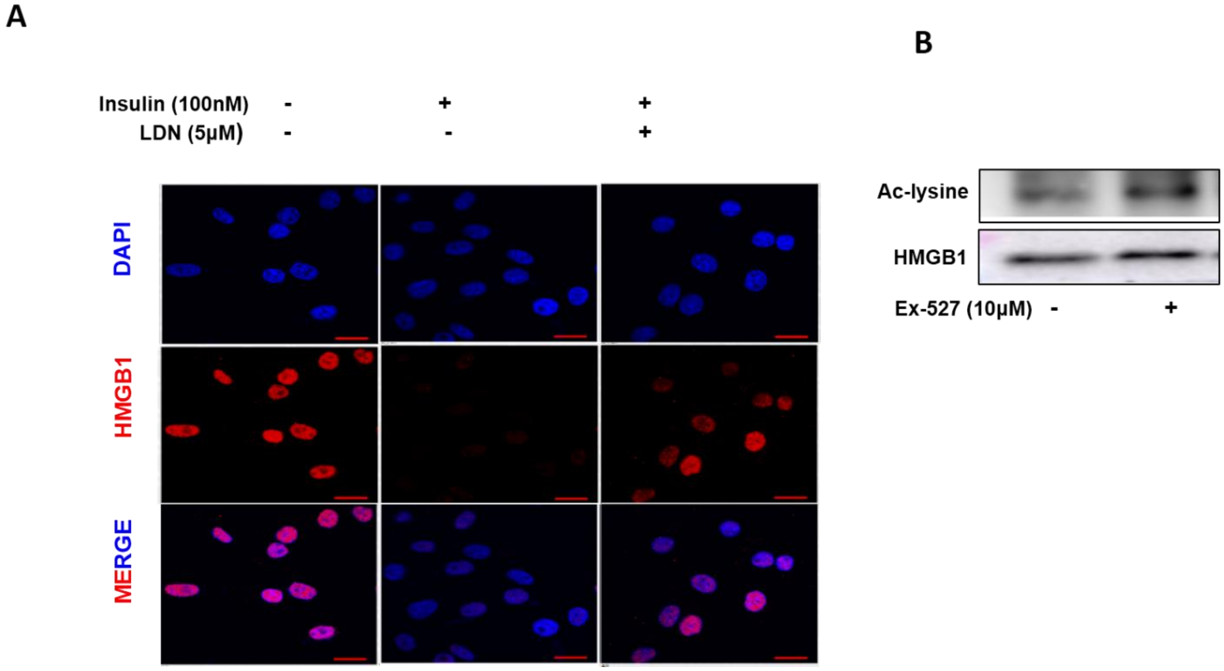


**Figure S3: SIRT1 regulates acetylation level of HMGB1: (A)** Immunocytochemistry for HMGB1 in HepG2 cells after 100nM insulin treatment in presence and absence of LDN for 4 hrs. showing 40X magnification, pseudocolouring: red: HMGB1,blue: nucleus counter-stain with DAPI . **(B)** Macrophage cells were incubated with or without 10µM Ex-527 (SIRT-1 inhibitor) and cell supernatant were concentrated using Amicon filter and immunoprecipitated with anti-Flag and IB with acetylated lysine and HMGB1. level of acetylated HMGB1 in immunoprecipitates increased significantly in presence of Ex-527 treatment.

**SIRT1 activity assay:**

Naltrexone SIRT1 deacetylase activity was measured using SIRT1 assay kit (CS1040, Sigma-Aldrich, USA) according to the manufacturer's instructions. Briefly1 µg of SIRT1 in combination with or without LDN (5µM) was incubated for 10 min and followed the protocol mentioned in SIRT1 assay kit. Fluorescence intensity were measured at Excitation370nm and Emission 460 nm (Infinite M200 Pro TECAN). Values were represented as fold change with respect to control.

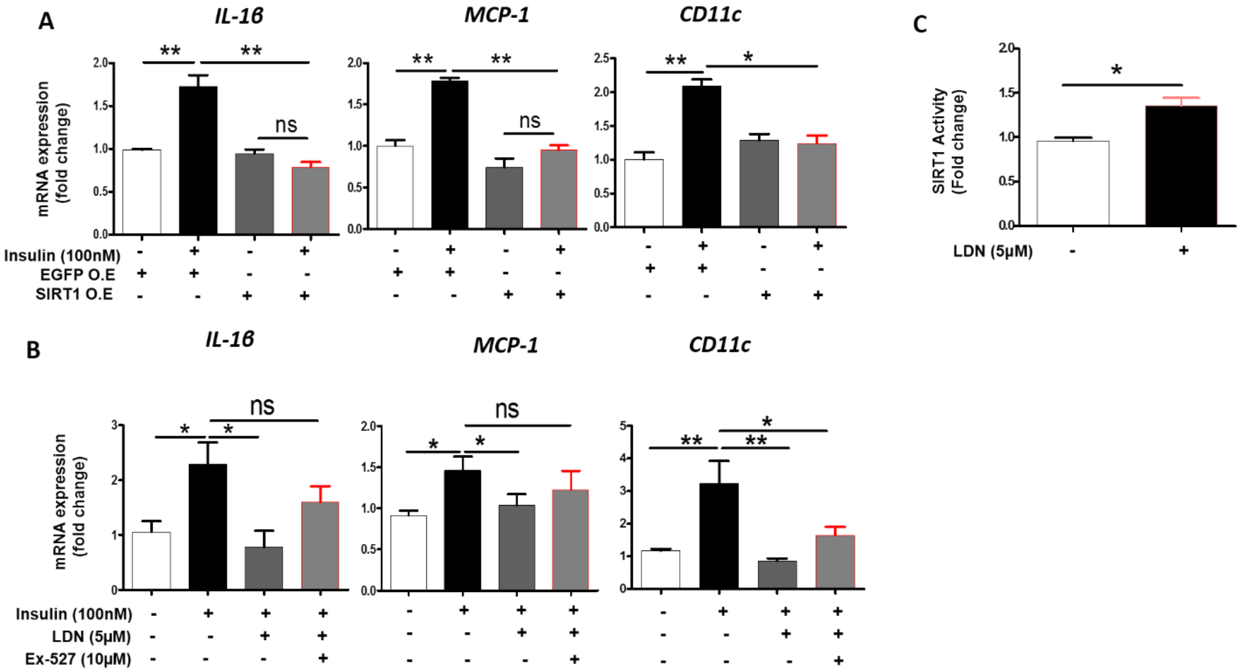


**Figure S4: (A)** Macrophage cells were transfected with SIRT1 and EGFP plasmid, incubated with or without 100 nM insulin for 24 hrs and mRNA levels of inflammatory genes were analyzed**. (B)** Macrophage cells were incubated with 100 nM insulin and LDN in presence and absence of 10µM EX-527 and mRNA levels of inflammatory genes were analyzed. **(C)** Naltrexone SIRT1 deacetylase activity, fluorescence intensity was measured at excitation370nm and emission 460 nm. Values were represented as -fold change with respect to control (mean ± SEM *p<0.05.)

**Characteristics of subjects:**

Total 8 no of subjects were included in this study and classified into two groups: 1. Healthy volunteers (Control), 2. Type-2 diabetic (Diabetic).

**Inclusion:** The subjects between 30 to 40 years of age were included in our study. 1. Healthy volunteer. 2. Type-2 Diabetic - newly diagnosed and treatment naive. Exclusion**:** BMI over 25, female, hypertension, lunatic and subjects with any other comorbidity were excluded.

**1. Healthy volunteer:**

| S.no | Age | Sex | BMI | Fasting glucose(mg/dL) | HbA1c (%) | Fasting insulin (µU/mL) | C.peptide (ng/mL) | HMGB1 (ng/mL) |
|---|---|---|---|---|---|---|---|---|
| 1 | 36 | M | 24.9 | 87 | 5.6 | 4.94 | 1.32 | 2.327 |
| 2 | 40 | M | 24.8 | 64 | 5.2 | 7.12 | 1.52 | 2.386 |
| 3 | 34 | M | 23.9 | 88 | 5.7 | 5.13 | 1.55 | 3.134 |
| 4 | 38 | M | 24.7 | 76 | 5 | 4.66 | 2.03 | 2.470 |

**2. Type-2 diabetic:**

| S.no | Age | Sex | BMI | Fasting glucose(mg/dL) | HbA1c(%) | Fasting insulin (µU/mL) | C.peptide (ng/mL) | HMGB1 (ng/mL) |
|---|---|---|---|---|---|---|---|---|
| 1 | 40 | M | 24.6 | 140 | 6.1 | 9.09 | 3.21 | 4.02 |
| 2 | 38 | M | 24.5 | 160 | 8.9 | 12.5 | 2.51 | 4.59 |
| 3 | 39 | M | 24.7 | 190 | 12.4 | 14.12 | 3.29 | 5.84 |
| 4 | 38 | M | 24.8 | 254 | 13.5 | 14.8 | 4.06 | 5.75 |

**Table1: Antibody list**

| S.No. | **Antibody** | **Species-specific** | **Company** | **Cat No.** |
|---|---|---|---|---|
| 1. | Phospho-Akt (Ser473) | Rabbit | CST | 4058 |
| 2. | Akt | Rabbit | CST | 9272 |
| 3. | SIRT1 | Mouse | CST | 2028 |
| 4. | β-actin | Rabbit | CST | 4970 |
| 5. | β-actin | Mouse | SANTRCRUZ | sc-47778 |
| 6. | HMGB1 | Mouse | ABNOVA | H00003146-M08 |
| 7. | Phospho- NF-κB P65(ser536) | Rabbit | CST | 3033 |
| 8. | NF-κB P65 | Mouse | CST | 6956 |
| 9. | Anti-FLAG | Mouse | SIGMA | F1804 |
| 10. | Anti-rabbit (secondary antibody) | Rabbit | CST | 7074 |
| 11. | Anti-mouse (secondary antibody) | Mouse | CST | 7076 |
| 12. | Alexa Fluor® 647 donkey anti-mouse(IgG H+L) | Mouse | JACSON | 715-605-150 |
| 13. | Anti-tubulin | Mouse | THERMO-SCIENTIFIC | 62204 |
| 14. | Lamin A/C | Mouse | SANTRCRUZ | sc-7293 |
| 15. | IgG | Rabbit | SIGMA | I5006 |

**Table 2: Primer sequences**

| S.No. | **Gene** | **Forward primer (5'-3')** | **Reverse primer (3'-5')** |
|---|---|---|---|
| 1. | Mcp-1 | GAAGGAATGGGTCCAGACAT | ACGGGTCAACTTCACATTCA |
| 2. | IL-1B | CACAGCAGCACATCAACAAG | GTGCTCATGTCCTCATCCTG |
| 3. | CD11c | ATGGAGCCTCAAGACAGGAC | GGATCTGGGATGCTGAAATC |
| 4. | TLR-4 | CAATCGCATAGAGACATA | GTTCAACATTCACCAAGA |
| 5. | TNF-a | TCTTCTCATTCCTGCTTGTGG | GGTCTGGGCCATAGAACTGA |
| 6. | IL-6 | CTCTGGGAAATCGTGGAAAT | CCAGTTTGGTAGCATCCATC |
| 7 | IL-10 | ATAACTGCACCCACTTCCCA | GGGCATCACTTCTACCAGGT |
| 5. | ARG-1 | TTTTTCCAGCAGACCAGCTT | AGAGATTATCGGAGCGCCTT |
| 6. | CD68 | TTGCTAGGACCGCTTATA | AAGGATGGCAGGAGAGTA |
| 7. | 18-S | GCAATTATTCCCCATGAACG | GGCCTCACTAAACCATCCAA |
| 8. | TLR4-siRNA | GAGCCGUUGGUGUAUCUUUTT | AAAGAUACACCAACGGCUCTG |
